# Critical Differential Expression Assessment for Individual Bulk RNA-Seq Projects

**DOI:** 10.1101/2024.02.10.579728

**Authors:** Charles D. Warden, Xiwei Wu

**Affiliations:** Integrative Genomics Core, Department of Molecular and Cellular Biology, City of Hope National Medical Center, Duarte, CA

## Abstract

Finding the right balance of quality and quantity can be important, and it is essential that project quality does not drop below the level where important main conclusions are missed or misstated. We use knock-out and over-expression studies as a simplification to test recovery of a known causal gene in RNA-Seq cell line experiments. When single-end RNA-Seq reads are aligned with STAR and quantified with htseq-count, we found potential value in testing the use of the Generalized Linear Model (GLM) implementation of edgeR with robust dispersion estimation more frequently for either single-variate or multi-variate 2-group comparisons (with the possibility of defining criteria less stringent than |fold-change| > 1.5 and FDR < 0.05). When considering a limited number of patient sample comparisons with larger sample size, there might be some decreased variability between methods (except for DESeq1). However, at the same time, the ranking of the gene identified using immunohistochemistry (for ER/PR/HER2 in breast cancer samples from The Cancer Genome Atlas) showed as possible shift in performance compared to the cell line comparisons, potentially highlighting utility for standard statistical tests and/or limma-based analysis with larger sample sizes. If this continues to be true in additional studies and comparisons, then that could be consistent with the possibility that it may be important to allocate time for potential methods troubleshooting for genomics projects.

Analysis of public data presented in this study does not consider all experimental designs, and presentation of downstream analysis is limited. So, any estimate from this simplification would be an underestimation of the true need for some methods testing for every project. Additionally, this set of independent cell line experiments has a limitation in being able to determine the frequency of missing a highly important gene if the problem is rare (such as 10% or lower). For example, if there was an assumption that only one method can be tested for “initial” analysis, then it is not completely clear to the extent that using edgeR-robust might perform better than DESeq2 in the cell line experiments.

Importantly, we do not wish to cause undue concern, and we believe that it should often be possible to define a gene expression differential expression workflow that is suitable for some purposes for many samples. Nevertheless, at the same time, we provide a variety of measures that we believe emphasize the need to critically assess every individual project and maximize confidence in published results.

## Introduction

Several bulk RNA-Seq differential expression benchmarks have been previously published [1–8]. However, if data analysis is performed for a sufficient number of comparisons, then we expect that some methods troubleshooting may be needed. This might be consistent with a previously published concept of “post-differential analysis sanity checks” [9]. In addition to the code that is the basis for this publication, there is also independent publications showing value in giving users the ability to compare multiple methods for a given comparison [10–13]. Additionally, evidence from Williams et al. [4] supports the idea that a tradeoff between methods exists, even as a best-case-scenario. Among the comparisons in this manuscript, currently popular methods include DESeq2 [14], edgeR [15–18], and limma-voom [19].

However, we have not previously attempted to quantify the idea problems can occur if you are forced to use a single method for all bulk RNA-Seq comparisons. Additionally, examples using public data may be helpful to communicate experiences in terms of a general concept. There has been some precedence in showing that bulk RNA pre-processing steps can have some effect on the results [20], and we test some variation in preprocessing for some cell line datasets (with TopHat2 [21] + htseq-count [22], STAR [23] + htseq-count [22], and Salmon [24]). However, the primary emphasis of this manuscript is to describe variability in differential expression results, with the variability in a subset of preprocessing strategies providing some reference for context.

An intuitive example is to use knock-out or over-expression cell line experiments, where the causal gene is known. This does not cover all possible complications with bulk RNA-Seq differential expression analysis. However, if we can show that it can be problematic to lock down a single method to identify the causal gene in this clear example, then we believe that would provide precedent to quantify the claim that is based upon less formal communication of experiences.

We also use protein immunochemistry from TCGA breast cancer data [25], but the causal gene is less clear in that situation. Nevertheless, if an approximate sense of how often the number of genes identified can greatly vary, then we believe that compliments analysis of recovering causal genes for gene expression changes.

## Materials and Methods

### Overview of Cell Line Experimental Design

To make results more comparable to between studies (as well as 51 bp single-end HiSeq2500 RNA-Seq data previously generated by the Integrative Genomics Core), only the forward read was considered for all re-processed raw data. However, for either paired-end or single-end libraries, only the forward .fastq.gz reads were downloaded using ‘*wget*’ for the following datasets: E-MTAB-1994 (HIF1A knock-down, [26]), E-MTAB-2128 (AGO2 knock-down, [27]), E-MTAB-2277 (UNK knock-down, [28]), E-MTAB-2682 (U2AF1 knock-down, 2 siRNA designs, 2 library types, [29, 30]), E-MTAB-3021 (PRPF8 knock-down, [31, 32]), E-MTAB-3740 (FOXP1 knock-down, 2 cell lines), E-MTAB-3849 (DAZL+NANOS3 over-expression, [33]), E-MTAB-4009 (HNRNPC knock-down, [34]), E-MTAB-4091 (DDX6 knock-down, 2 cell fractions, [35]), E-MTAB-4237 (BMI1 knock-down, 2 cell lines), E-MTAB-5162 (RNH1 knock-down, [36]), E-MTAB-5566 (PBRM1 knock-down, 2 batches), E-MTAB-5577 (PATL1 knock-down, [35]), E-MTAB-6204 (MATR3 knock-down, 2 cell fractions, [37]), E-MTAB-6235 (DDX3X+DDX54 knock-down), E-MTAB-6264 (NFE2L2+NUAK1+RNPS1 knock-down, [38]), E-MTAB-6756 (GATA6+HNF4A knock-down, [39]), E-MTAB-7033 (TP63 knock-down, 2 cell lines, [40]), E-MTAB-7284 (TIGAR knock-down, [41]), E-MTAB-7791 (ARID1A siRNA + ARID1A CRIPSR, [42]).

After the data was downloaded, one of the following processing strategies was used: TopHat2 [21] alignment + htseq-count [22] quantification, STAR [23] alignment + htseq-count [22] quantification, or Salmon [24] quantification. When STAR was used as the aligner, the parameters “*--twopassMode Basic --outSAMstrandField intronMotif*” were also used. For genomic alignments, Read counts were quantified using htseq-count [22], with UCSC known gene annotations (TxDb.Hsapiens.UCSC. hg19.knownGene, [43]). GENCODE (v32, [44]) transcripts were used for transcriptome quantifications. For genomic alignments, aligned reads were counted using GenomicRanges [45]. This follows processing strategies based upon https://github.com/cwarden45/RNAseq_templates. The strandedness of the libraries was determined or checked using RSeQC [46], along with other tasks from *run_RSeQC.py* in the template.

A larger number of possible over-expression and/or knock-down experiments is listed in *Tentative_Datasets_to_Download.xlsx* on the SourceForge page (https://sourceforge.net/projects/rnaseq-deg-methodlimit). However, in the interests of time with this amount of manual work for each dataset, only the datasets deposited in the ENA (with accessions that start with “E-MTAB-”) that could be individually confirmed to have an experimental design consistent with the goals of this study were downloaded.

**Table 1:**
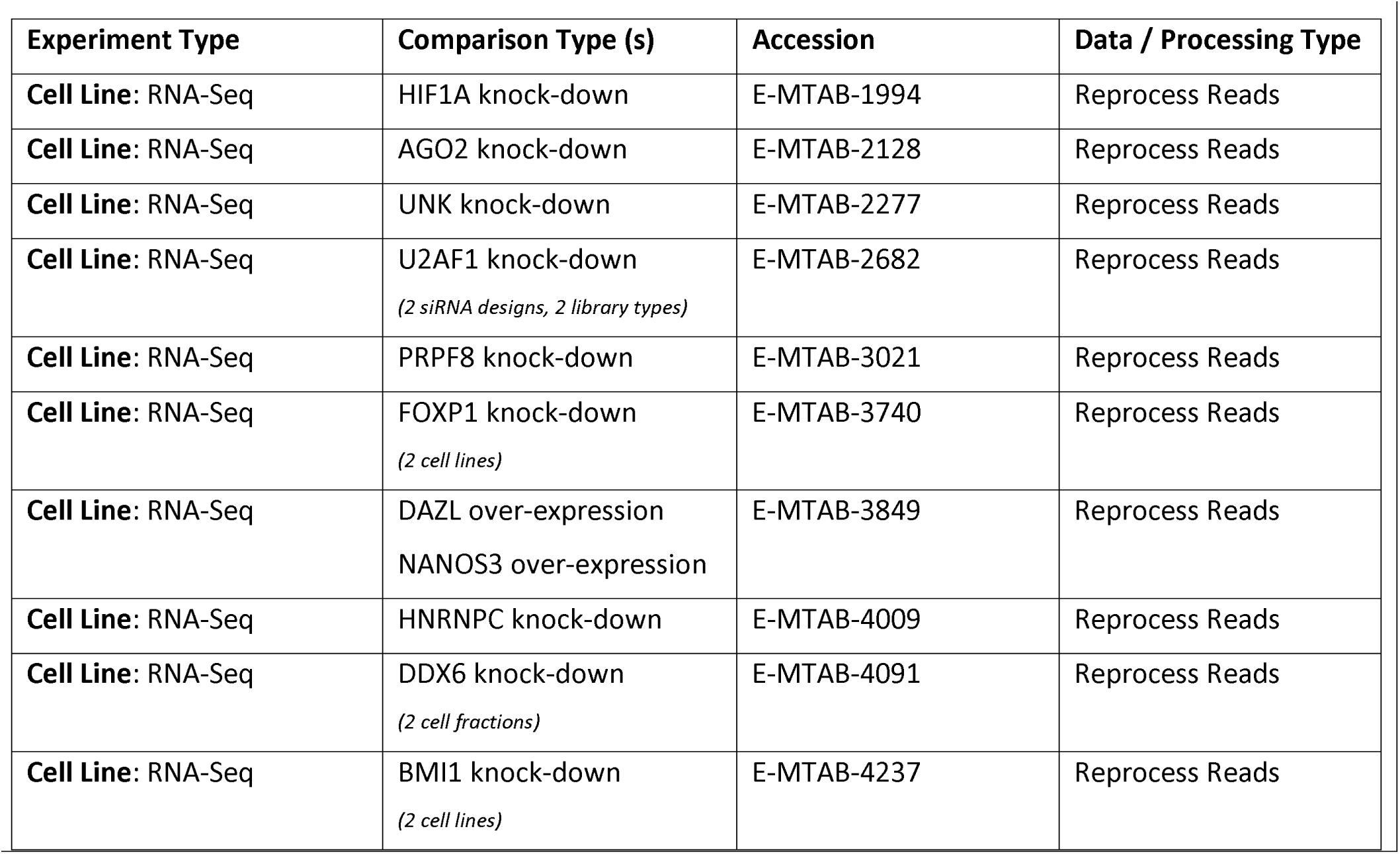
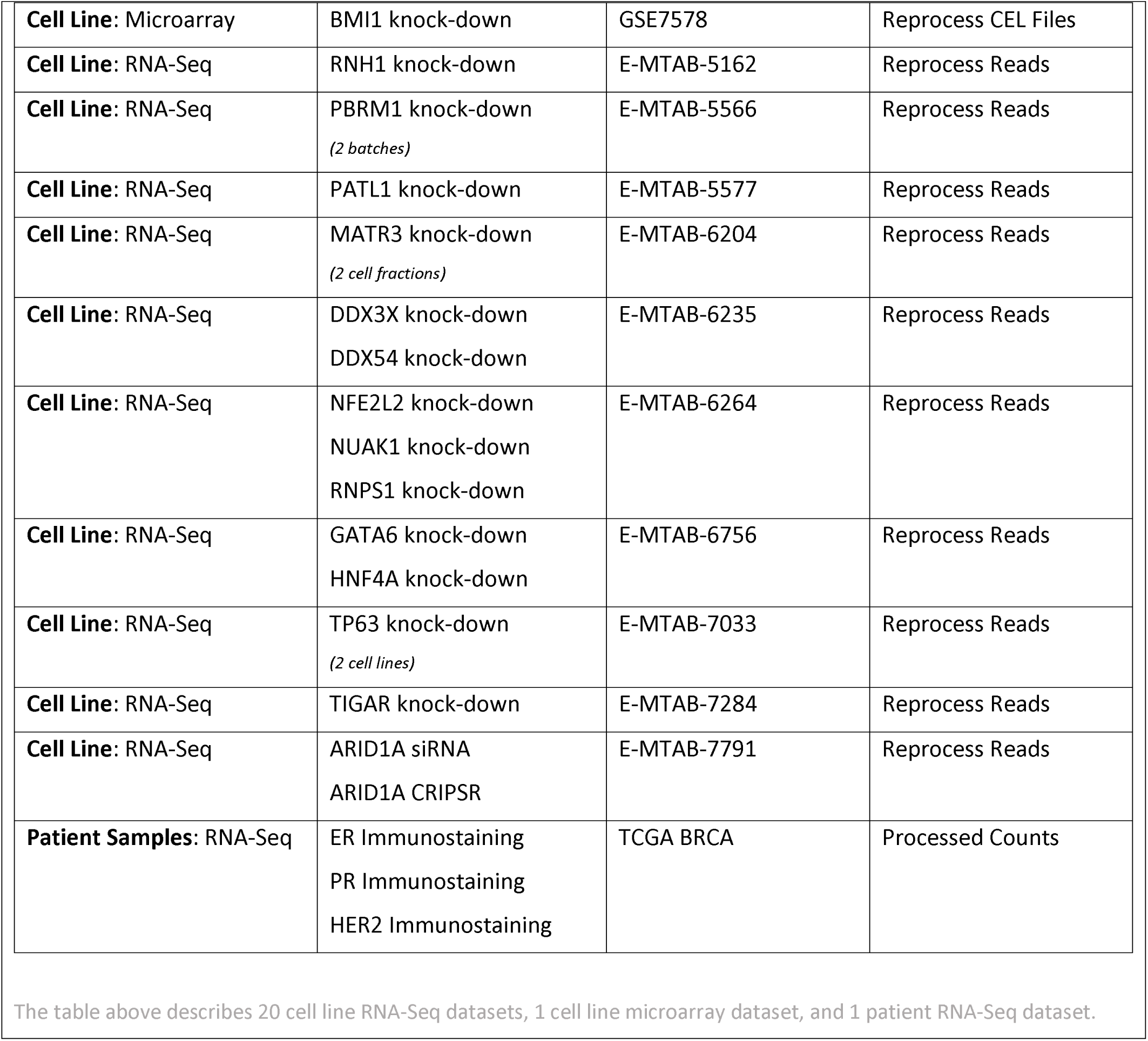
List of Public Datasets for Critical Assessment

### Comparisons of Differential Expression Implementations within R Packages

With or without the upstream processing steps, samples were processed and read counts were generated with strategies similar to presented in https://github.com/cwarden45/RNAseq_templates. Processing upstream of differential expression analysis in R has been briefly described in the *Overview of Cell Line Experimental Design* section. While not presented in the main paper, the .zip files including the full code and results also include some quality control (QC) plots for each separate study. Those files are saved in https://zenodo.org/records/3378055. Suggested QC plots that might facilitate evaluation of a dataset include dendrogram with hierarchical clustering, log2(FPKM/TPM + 0.1) box plots, Principal Components Analysis (PCA), and log2(FPKM/TPM + 0.1) density plots.

Fold-change values were calculated from Fragments Per Kilobase per Million reads (FPKM, [47]) normalized expression values, which were also used for visualization (following a log_2_ transformation). Using log2(FPKM + 0.1) is an intuitive way to use limma-trend, given that also used for fold-change value that is usually calculated independently of the p-value calculation. When using limma-trend with Count-Per-Million (CPM) expression, the *cpm()* function from edgeR was used with the “*log=TRUE, prior.count=3*” parameters (matching [48] as well as the limma user guide). While not included in the main manuscript, there are heatmaps and volcano plots generated using log2(FPKM + 0.1) values within the separate .zip files with code and results for each study (described above). When heatmaps were generated, they either used the *heatmap.2* function (from the ‘*gplots*’ R package) or the *heatmap.3* function (downloaded from https://github.com/obigriffith/biostar-tutorials/blob/master/Heatmaps/heatmap.3.R).

Prior to p-value calculation, genes were filtered to only include transcripts with an FPKM expression level of 0.1 (after a rounded log2-transformation) in at least 50% of samples [49] as well as genes that are greater than 150 bp. For all comparisons, the following strategies were compared: *i)* edgeR Generalized Linear Model (edgeR-GLM), *ii)* edgeR Generalized Linear Model with robust dispersion estimate (edgeR-GLM-robust), *iii)* edgeR Quasi-Likelihood (edgeR-QL), *iv)* edgeR Quasi-Likelihood with robust dispersion estimate (edgeR-QL-robust), *v)* DESeq1, *vi)* DESeq2, *vii)* limma-voom, *viii)* limma-trend using edgeR CPM values (limma-trend-CPM), *ix)* limma-trend using log2(FPKM/TPM + 0.1) values (limma-trend-FPKM.TPM), and), *x)* ANOVA using log2(FPKM/TPM + 0.1) values (ANOVA). False discovery rate (FDR) values were calculated using the method of Benjamini and Hochberg (B-H FDR, [50]). At least for initial settings, genes were defined as differentially expressed if they had a |fold-change| > 1.5 and FDR < 0.05.

In general, there are multiple ways to run edgeR. For either edgeR-GLM-robust or edgeR-QL-robust, *estimateGLMRobustDisp()* is used to estimate dispersion values, but Lun et al. 2016 [51] indicate that this not affect downstream analysis for the QL method. So, our expectation is that differences for the edgeR-QL-robust strategy should be due to using “*robust=TRUE*” parameter for *glmQLFit()*, which was part of the workflow for Lun et al. [51] (referencing [52] for the robust QL estimation). Also, the QL strategy was tested at a later point in time: so, please note that the overall strategy called “*edgeR-GLM*” uses *estimateCommonDisp()* for dispersion estimates, whereas the overall strategy called “*edgeR-QL*” uses *estimateDisp()* for dispersion estimates.

Cuffdiff [53] was also run when a relevant 2-group comparison could be defined. However, the causal gene was usually not recovered when run with the configuration in this study. This may be similar to some previous benchmark publications [1, 54]. So, summary statistics are provided in *Target_Recovery_Status.xlsx* on the SourceForge page (https://sourceforge.net/projects/rnaseq-deg-methodlimit). However, cuffdiff is not presented among the main results or Supplemental Tables in this manuscript. It is not currently clear if only using the forward read for re-alignment in this study might have decreased the performance of cuffdiff, although some mention of decreased performance has been previously reported [1].

### Overview of ER/PR/HER2 Subtype Analysis in TCGA BRCA Data

Unlike cell line data, raw reads were not reprocessed for samples from The Cancer Genome Atlas (TCGA), in part due to being controlled access and in part to try and use some limitations in total computation for this particular study. Unstranded counts and unstranded FPKM values were downloaded using TCGAbiolinks [55]. The full script used to download and reformat data is contained within https://zenodo.org/records/3378055/files/TCGA_BRCA.zip.

Modified templates for the RNA-Seq analysis used for cell line analysis are also included in the above link. Some general changes will briefly be provided here. Under an assumption of greater heterogeneity for patient data, the criteria for differential expression was |fold-change| > 1.2 and FDR < 0.05 (instead of |fold-change| > 1.5). In addition to the cell line method comparisons, the non-parametric Wilcoxon rank-sum test was also compared (as recommended by [56]). The expression criteria for genes to test for differential expression were also varied more than the cell line comparisons, as described in the Results section. Unlike the cell line comparisons, the FPKM values provided by TCGAbiolinks were used and FPKM values were not re-calculated from counts (where the rounding factor for calculating the fold-changes also no longer varies with the expression criteria). There were also various smaller changes made to the code, possibly similar to changes needed to modify the RNA-Seq analysis template for different cell line studies.

### Testing Additional Alignment and Quantification for Selected Datasets

The primary goal of this manuscript is to compare differential expression for bulk RNA-Seq data. As such, we provide some variation in preprocessing for comparison. However, we do not provide testing for all possible popular workflows. For a couple datasets (E-MTAB-2128 and E-MTAB-3740) where the causal gene is never identified with |fold-change| > 1.5 and FDR < 0.05, we also provide some additional testing with HISAT2 [57] and kallisto [58]. These were selected on the basis of having complications with all of the 3 main processing strategies used for cell line data, and summary statistics are included in **Supplemental Table S1**.

Limited transcriptome alignments followed by quantification were also performed for E-MTAB-2128 and E-MTAB-3740. Namely, Bowtie1 [59] alignments were further processed for quantifications using RSEM [60], and Bowtie2 [61] alignments were further processed for quantifications using eXpress [62]. Dot plots for the causal genes (AGO2 and FOXP1, respectively) are shown in **Supplemental Figure S1** and **Supplemental Figure S2**. Qualitatively, the Bowtie1+RSEM quantification may be more similar to the genomic alignments for AGO2 (E-MTAB-2128) but more similar to Salmon/kalliso for FOXP1 (E-MTAB-3740).

As an alternative to htseq-count, featureCounts ([63], from the subread package) was also tested for E-MTAB-2128 and E-MTAB-3740 using the original STAR alignments. Slightly different counts of differentially expressed genes were identified when using featureCounts versus htseq-count (**Supplemental Table S1**). Additionally, fastp [64] was used to test upstream read filtering with a STAR alignment of filtered and trimmed reads (with parameters “*--length_required 40 --n_base_limit 0*”). Upstream application of fastp had minimal effect on E-MTAB-3740, but upstream application of fastp had a noticeable effect on starting read count for E-MTAB-2128 that was influenced by a higher frequency of degenerate nucleotides in the reads (**Supplemental Table S1**). Gene counts not identical for either E-MTAB-2128 orE-MTAB-3740, but fastp did not help with identifying the causal gene for E-MTAB-2128.

Newer samples are more likely to be processed with hg38 than hg19, and newer samples may be paired-end instead of single-end. So, as a partial test for E-MTAB-2128 and E-MTAB-3740, the UCSC hg38 sequence was also downloaded from iGenomes. To be more comparable to the similar hg19 sequence, the supplemental chromosomes were removed to create a reference only using “canonical” chromosome sequences from the reference genome. Copies of the scripts used to accomplish this are provided in the “*Code/Round12*” subfolder for the .zip file with additional results and full code for Round 12 for E-MTAB-2128. An index for the STAR alignments was then created. Similarly, a UCSC Known Gene GTF file was used for genomic annotation using the *TxDb.Hsapiens.UCSC.hg38.knownGene* Bioconductor package, with annotations downloaded as available on 11/24/2022. Pre-defined housekeeping gene annotations were downloaded for hg38 as provided by RSeQC (https://sourceforge.net/projects/rseqc/files/BED/Human_Homo_sapiens/). For both E-MTAB-2128 and E-MTAB-3740, switching to use hg38 improved the alignment rate more than running fastp upstream of a STAR alignment.

Finally, the use of a different annotation strategy for the genome alignment and transcriptome quantification adds a confounding factor to compare preprocessing. So, as a partial attempt to consider the gene annotation source, *gffread* ([65], from the cufflinks package) was used to convert the UCSC Known Gene annotations from Bioconductor to a FASTA transcriptome file. The original GTF did not contain transcriptome annotations, so “gene_id” was replaced with “transcript_id” before running *gffread*. The resulting FASTA file was indexed with Salmon (with the same settings as the GENCODE transcriptome reference), and performance was tested for the 3 comparisons from E-MTAB-2682.

### Gene Enrichment Analysis

A limited set of gene sets were tested for enrichment, based upon the causal gene affected in certain experiments (BMI1 from E-MTAB-4237, HIF1A from E-MTAB-1994, and GATA6 from E-MTAB-6756). There are “UP” and “DN” gene sets that exists within both c2 and c6 in MSigDB for BMI1, and there similar c2 signatures for HIF1A, and GATA6. If using upstream differentially expressed gene lists / statistics, then the DESeq2 and edgeR-robust-GLM results were used. Analysis for c6 gene sets (but not c2 gene sets) can be performed using Enrichr. GSEA and BD-Func analysis can be performed with both c2 and c6 gene lists and any pair of “UP” and “DN” c2 or c6 gene lists, respectively.

For Enrichr [66], only the MSigDB c6 signatures were available to test (for genes from DESeq2 and edgeR-robust-GLM differentially expressed gene lists, defined with |fold-change| > 1.5 and FDR < 0.05) for “*MSigDB Oncogenic Signatures*” on 12/25/2022 (under “*Diseases/Drugs*”). So, Enrichr was only used to test enrichment for BMI1. As long as there were at least triplicates per group, all 3 gene sets were tested with GSEA (v4.3.2, [67, 68]) using *i)* provided log2(FPKM + 0.1) expression values and *ii)* using a signed pre-ranked value defined as -log10(p-value) for positive fold changes and log10(p-value) for negative fold changes (for DESeq2 and edgeR-robust-GLM). For GSEA, Molecular Signatures Database (MSigDB, [69]) v2022.1 gene symbol signature files were used. Additionally, BD-Func (v1.1.7, [70]) was used to test enrichment, with an updated set of pairs of MSigDB c2 signatures for this manuscript. One method of applying BD-Func was previously described: scores were defined based upon log2(FPKM + 0.1) expression values for individual samples and an additional test on those scores was performed outside of BD-Func (such as a 1-way ANOVA or 2-way ANOVA). Additionally, BD-Func has a mode to compare pairs of up-regulated versus down-regulated gene lists for fold-change values: so, that strategy was modified to compare signed -log10(p-value) scores instead of the fold-change values calculated independent of the p-value calculation (for DESeq2 and edgeR-robust-GLM). The associated code is available in the associated .zip files for Round 8 of E-MTAB-1994, E-MTAB-4237, and E-MTAB-6756.

The c6 MSigDB signatures considered for significance and ranking are BMI1_DN.V1_UP [71] and BMI1_DN.V1_DN [71]. The c2 MSigDB signatures considered for significance and ranking are DOUGLAS_BMI1_TARGETS_UP [72], DOUGLAS_BMI1_TARGETS_DN [72], ELVIDGE_HIF1A_TARGETS_UP [73], ELVIDGE_HIF1A_TARGETS_DN [73], ELVIDGE_HIF1A_AND_HIF2A_TARGETS_UP [73], ELVIDGE_HIF1A_AND_HIF2A_TARGETS_DN [73], ZHANG_GATA6_TARGETS_UP [74], and ZHANG_GATA6_TARGETS_DN [74].

### Microarray Re-Analysis

For one casual gene (BMI1), the original data for a surprising MSigDB signature was checked (for BMI1_DN.V1_UP and BMI1_DN.V1_DN [71]). Code for analysis is based upon https://github.com/cwarden45/HuGene_Expression_Template. Affy Power Tools (v2.11.6) was used to define a custom gene symbol summarization to use or differential expression (with |fold-change| > 1.5, limma [75] p-value < 0.05, B-H FDR [50] < 0.05, for genes with 50% RMA expression > 3.5).

Because shBMI1 was used as the treatment group for the microarray experiment, the direction should be reversed relative to regular BMI1 regulation. However, the RNA-Seq experiment from E-MTAB-4237 is also a knock-down experiment. So, similar to the MSigDB signatures, the direction of change is the opposite of what would be expected (even though all 3 BD-Func results were significant with an unadjusted p-value < 0.05, for a single test of a custom signature). Full code and results are provided in https://zenodo.org/records/3378055/files/GSE7578.zip. Application of the custom gene signature was applied in Round 9 of the analysis for E-MTAB-4237 (https://zenodo.org/records/3378055/files/E-MTAB-4237.zip).

### Code Availability

Links to available data are available from https://sourceforge.net/projects/rnaseq-deg-methodlimit/. This includes links to code and results to individual datasets, such as provided on Zenodo: https://zenodo.org/records/3378055.

### Example of Custom Code Modifications

One of the most common examples of manual modification is revision of figures in preparation for publication. This is true for the full set of code and results linked from this study (such as modifying labels in heatmaps, and the number of variables displayed that may sometimes need to be represented in multiple annotation tracks) as well as for other projects. However, from an analysis standpoint, this might be considered minor.

Within this study, another example is the need to modify the code to run DESeq1 for paired samples without replicates within each paired group. As mentioned in the next section, there are occasional brief notes in the public LOG (which can be searched for “DESeq1”). However, as a specific example, the design for E-MTAB-3740 required a modification relative to the other similar comparisons in order to run without generating an error message due to the lack of replicates within each group defined by the interaction of 2 variables used for the multivariate analysis (where “*dds = estimateDispersions(dds, method = “pooled”)*” needed to be changed to the equivalent of “*dds = estimateDispersions(dds, method = “pooled-CR”, modelFrame=colData, modelFormula=formula(count∼var1 + var2))*”). Strictly speaking, this could be accomplished in a pipeline with the addition of another variable and an additional case to define the code to use for differential expression. Nevertheless, this conceptually reflects something beyond a change that can be specified in a parameter file (such as choosing to run some particular configuration for “edgeR”, “DESeq2”, and/or “limma-voom”). Thus, the word “template” is used for repositories like https://github.com/cwarden45/RNAseq_templates, whereas a pipeline was code that doesn’t change for a given version (but there can be different configurations to optimize for different projects).

### Public Lab Notebook and Comments Regarding “P-hacking” and Methods Testing Misuse

When a need for methods testing for every project is emphasized, there is a risk that unrepresentative results could be selected to over-emphasize a point. This can be referred to as “p-hacking” [76].

We acknowledge that this is a risk. It may not be a complete solution, but we have attempted to create a basic log file (https://sourceforge.net/projects/rnaseq-deg-methodlimit/files/LOG.txt/download) to act like a public lab notebook. In this paper, we also go through all the details of testing analysis with different methods. The full details may create difficulties for communication of biological conclusions in a paper not centered on methods testing. However, if additional details are provided in a repository like GitHub and/or a public lab notebook, then we hope some record of all testing performed and the basis for selecting a fair and objective representative main result can help. The level of detail provided in this log file is limited, but additional details are provided in the summary slides within the “Results” subfolder of the compressed files with full code and results (https://zenodo.org/records/3378055).

## Results

### Noticeable Lack of Recovery for Casual Knock-Down or Over-Expressed Gene for Cell Line Experiments

There is some variation between different methods (including the mix of 3 preprocessing strategies), but no method can recover the causal gene for all studies and preprocessing strategies with |fold-change| > 1.5 and FDR < 0.05 (**Figure 1**, **Supplemental Table S1**).

**Figure 1:**
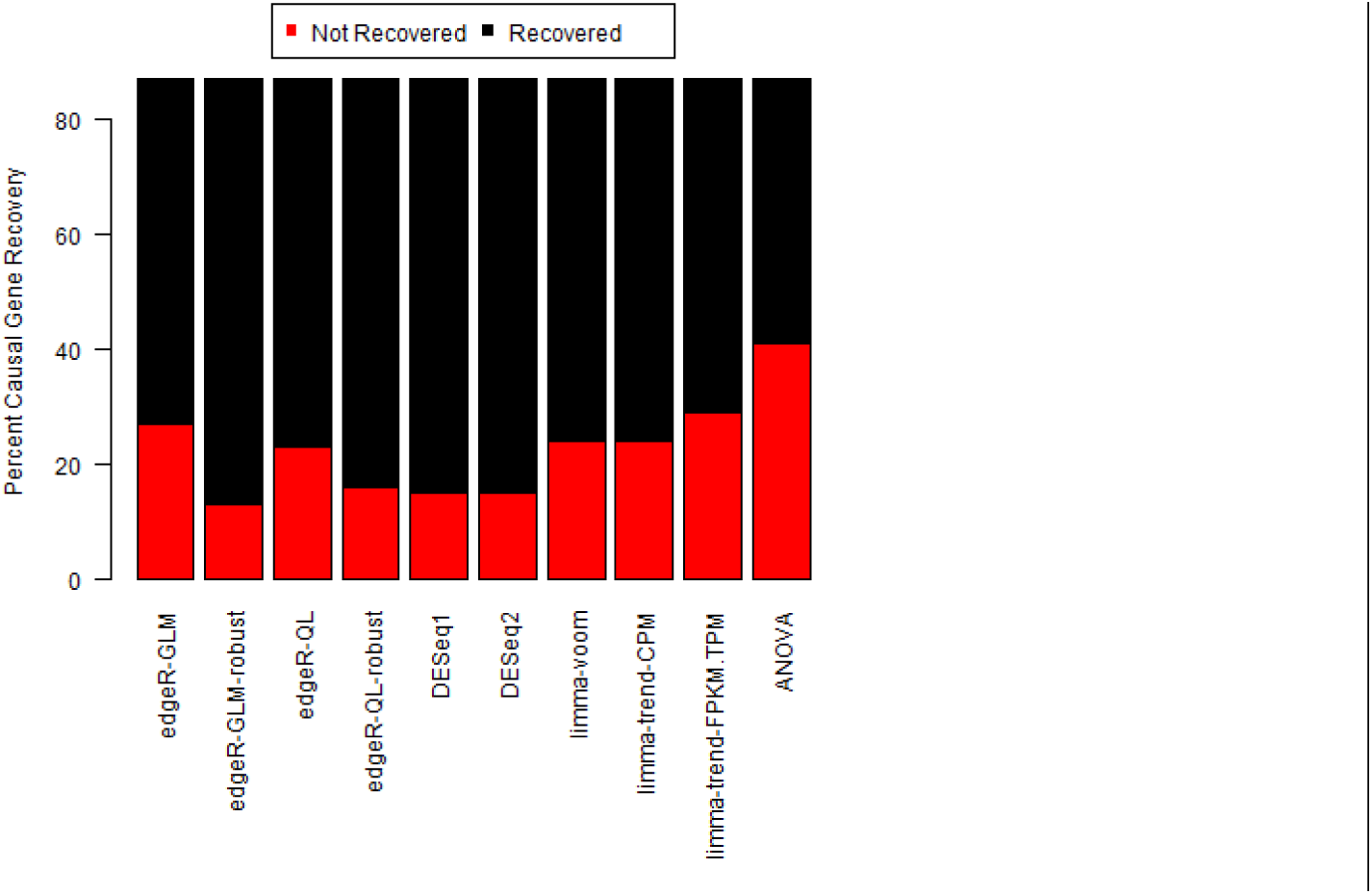
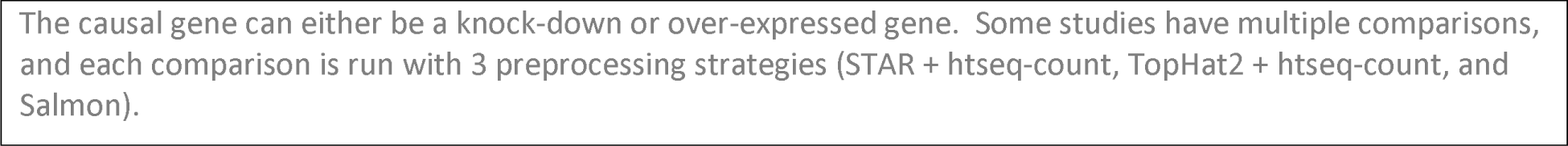
Recovery of causal gene with |fold-change| > 1.5 and FDR < 0.05.

If the preprocessing strategies are separated out and the FDR level is considered without a fold-change threshold, then there appears to be some advantages to using a quantification from a genomic alignment (**Figure 2**). There are other studies indicating advantages to pseudoalignment methods [77]. However, we believe that the results of this study are consistent with other published studies [78–80], or this at least matches the need to have a carefully considered set of decoy sequences [81].

**Figure 2:**
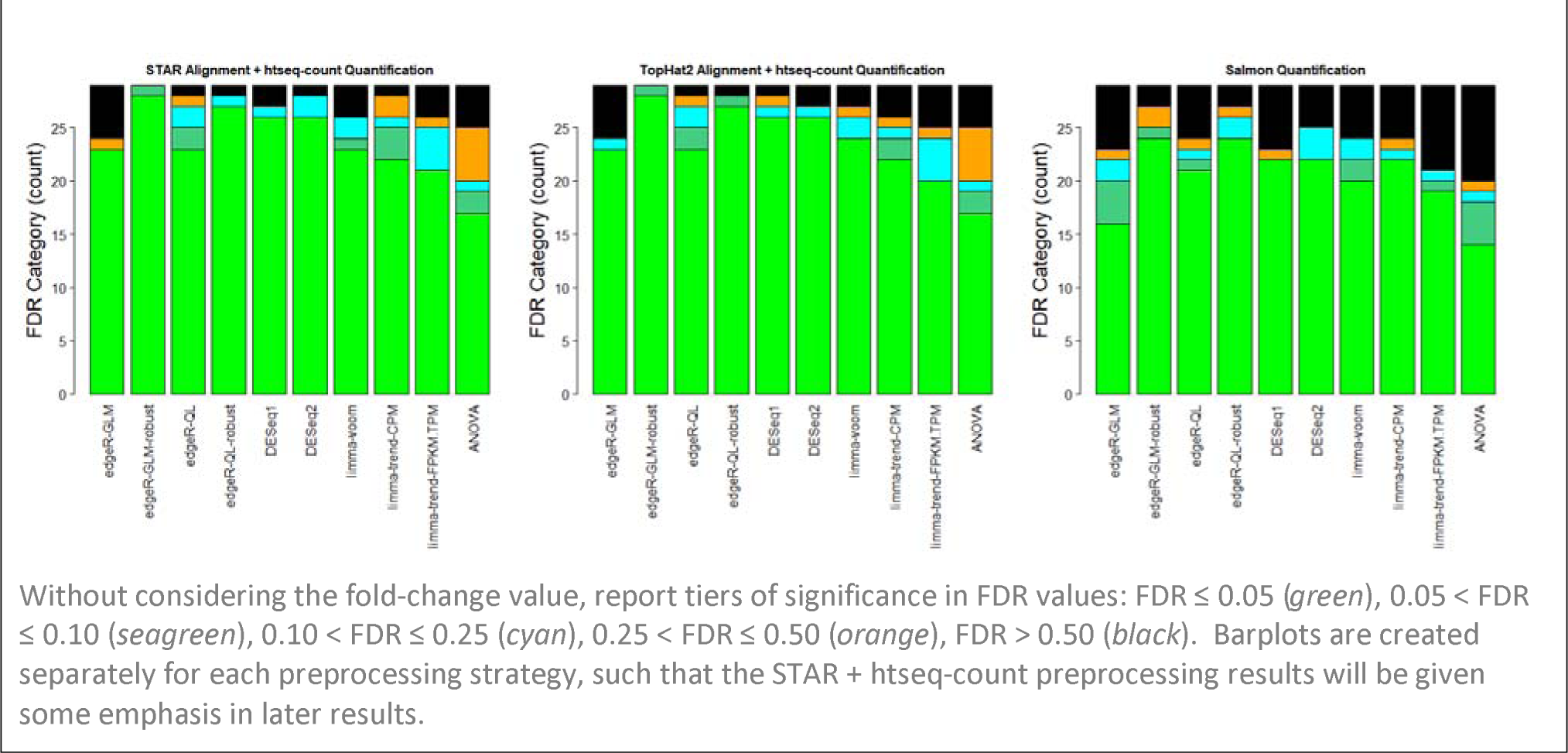
Some Evidence for Improvement for Causal Gene Recovery for STAR (and TopHat2)

For E-MTAB-2128 (and E-MTAB-3740), some additional variations in preprocessing were performed (*Supplemental Table S1* and **Supplemental Figure S1**). While some trends are different for genomic alignments versus transcriptomic quantifications, no tested strategy increases the fold-change for *AGO2* to be above 1.5 for in E-MTAB-2128. For E-MTAB-3740, edgeR-GLM-robust already has an FDR < 0.05 for the casual gene with the TopHat2 alignment (*FOXP1* in **Supplemental Figure S2**). The FDR value with that same method is marginally not significant at that threshold for the STAR and HISAT2 alignments. However, we do not recommend testing different alignments in order to change an FDR value to be marginally below 0.05, and we would consider that to be p-hacking. Instead, we would recommend increasing the FDR to identify more gene candidates (such as FDR < 0.10, FDR < 0.25, or possibly FDR < 0.50). Nevertheless, many significant results for the STAR alignments could be found with TopHat2 alignments, and there may occasionally be genuine benefits to using a TopHat2 alignment (such as possible increased stringency in alignments [82]). However, we do not consider the impact of unique files provided by TopHat2 or STAR in this study (such as for unaligned reads, gene fusions, splice junctions, etc.).

Given the result in **Figure 2**, the question could be raised if having a separate fold-change threshold is helpful (in that the causal gene can be lost when imposing these criteria: **Figure 1** and **Supplemental Table S1**). However, we believe that the fold-change calculated independently from the p-value calculation (and FDR calculation) provides useful information. For example, the worst ranking for the causal gene was better when ranking by fold-change rather than by p-value or FDR (**Supplemental Figure S3**). In absolute terms, the fold-change ranking was less than or equal to the p-value ranking for 366 out of 580 comparisons (63.1%). However, this was only a relatively consistent and large difference when using ANOVA for log2(FPKM + 0.1) expression values, and perhaps to some extent the GLM edgeR implementation. Importantly, a *signed* p-value ranking is created (using -log10(p-value) for positive fold-change values and log10(p-value) for negative fold-change values), then the fold-change ranking was less than or equal to the *signed* p-value ranking for 283 out of 580 comparisons (48.8%, **Supplemental Figure S4**). So, comparisons where ranking causal genes by fold-change was preferable are outliers, and gene ranking of genomic alterations by p-value should often be acceptable (for cell line experiments with two groups to compare). Additionally, the fold-change helps with interpretation of the p-value, and the fold-change was necessary to be able to define the signed p-value. Nevertheless, the ∼2% overall benefit to using a *signed* p-value was statistically significant by a Wilcoxon signed rank test (p=0.00011).

The use of the Bioconductor UCSC Known Gene annotations for the genome alignment and the GENCODE sequences for the transcriptome quantification adds a potential confounding factor in interpreting the causes of differences in results. At least for the annotation file created, exons were not defined with respect to transcripts for the UCSC Known Gene annotations. So, this is not perfectly transferable for a transcriptome quantification. Nevertheless, there was a noticeable decrease in performance in recovering the U2AF1 knock-down (among the 3 comparisons with E-MTAB-2682 data). There is improvement in recovering U2AF1 knock-down (**Supplemental Table S1**), when using a “transcriptome” based upon the gene-centric exon annotations. However, we do not generally recommend this as a strategy for running transcriptome quantification, since varying transcripts for the same gene are not well defined. Nevertheless, the use of annotations from a different source may help with the ability to identify the causal gene in this specific experiment.

### Variation in the Number of Differentially Expressed Genes for Cell Line Experiments

If we focus on the results for preprocessing that uses STAR for alignment and htseq-count for quantification, then we can see that the number of genes identified by method can vary (**Supplemental Figure S5**). For individual comparisons, adding the robust dispersion parameter estimation had a greater maximal effect for the Generalized Linear Model (GLM) than the Quasi-Likelihood (QL) comparison (**Supplemental Figure S6**).

We do not consider the amount of difference between edgeR-robust (GLM) and DESeq2 to be sufficient to conclude edgeR-robust should always be used over DESeq2 for differential expression. However, tentatively, we might consider the combination of edgeR-robust (GLM), DESeq2, and limma-voom to be sufficient as a starting point to guess what might be a good “initial” p-value calculation methods for comparisons for a particular project. Focusing on the public data analysis in this study, it may be helpful to focus on the extent of difference in results for edgeR-robust (GLM) versus DESeq2 (**Supplemental Figure S7**). When considering results based upon htseq-count from the STAR alignment, the number of genes identified by edgeR-robust (GLM) was significantly more than DESeq2 by Wilcoxon signed rank test (p=1.1 x 10). There is clearly more of a different compared to any other method, for either edgeR-robust (GLM) or DESeq2. However, to provide context, the number of genes for each method was compared to the median number of genes identified with any other method. For edgeR-robust (GLM), the number of genes showed a statistically significant increase compared to the median number of genes identified with any other method (p=7.5 x 10, Wilcoxon signed rank test). In contrast, the difference was not significant for DESeq2 versus the median number of genes identified with any other method (p=0.55, Wilcoxon signed rank test). Given that more alternative implementations for edgeR were among the “other” methods, we believe there is evidence that increased recovery of the causal gene may be due in part to a tendency to show a moderate increase in the number of genes identified.

Variation in the number of genes identified for a given study with different methods can also be observed (**Figure 3**). Please note that some comparisons have roughly similar number of genes identified with different methods. However, each individual comparison may be like the project for an individual lab. So, when the number of genes identified can vary by more than 3,000 genes, then only considering one particular method could be problematic.

**Figure 3:**
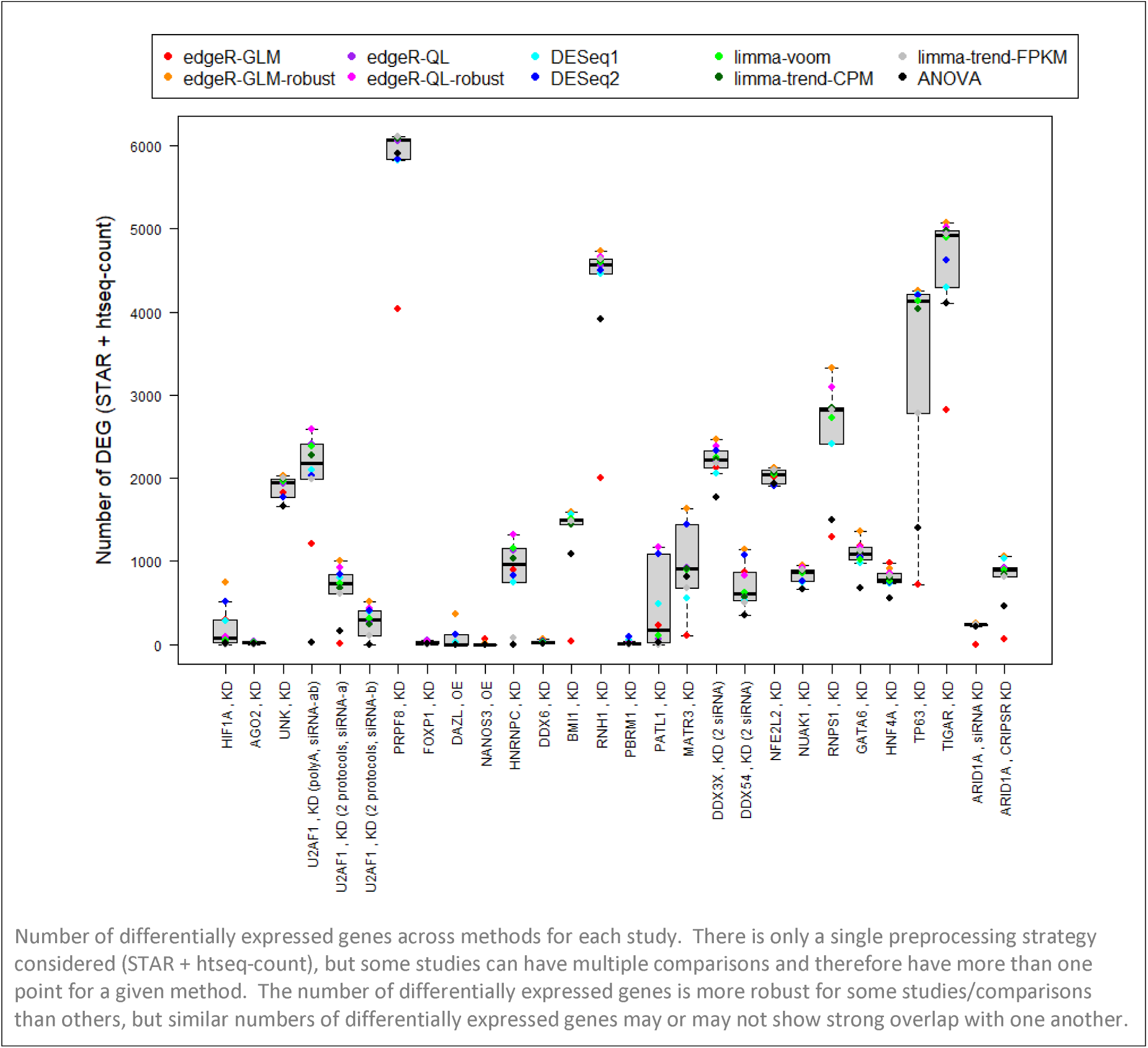
Number of Differentially Expressed Genes Across Methods by Cell Line Comparison

### Variation in the Number of Differentially Expressed Genes for A Larger Patient Cohort

While not precisely the causal gene, similar analysis can be performed for immunohistochemistry status breast cancer samples from The Cancer Genome Atlas project (TCGA BRCA, [25, 83]). In particular, the immunohistochemistry status was compared to the RNA-Seq mRNA differential expression status for estrogen receptor (ER, ESR1), progesterone receptor (PR, PGR), and human epidermal growth factor (HER2/ERBB2). In a previous study, there were some noticeable differences in the genes identified with different methods, even with varying sample sizes [49]. This was also observed in a more recent study by another group, but with an emphasis on using a non-parametric Wilcoxon rank-sum test for 2-group comparisons in larger sample size [56].

For the TCGA comparisons describe above, the provided data was analyzed with four tests of genes to test (with |fold-change| > 1.2 and FDR < 0.05): *i)* all genes with 50% FPKM > 0.1, *ii)* protein-coding genes with 50% FPKM > 0.1, *iii)* all genes with no upstream expression filter, *iv)* protein-coding genes with no upstream expression filter (**Supplemental Figure S8A**, **Supplemental Table S1**). Relative to the set of genes compared, the number of genes was largest for ER/ESR1 (n= 757), intermediate for PR/PGR (n= 754), and relatively lowest for HER2/ERBB2 (n= 548). The fold-change threshold was decreased for the patient data, but that might not have been needed in this situation, as most gene lists identified more than 2,000 differentially expressed genes. The effect of the upstream gene filter was relatively small compared to the immunohistochemistry status used for the comparison, and the direction in the change in number of genes identified also varied with comparison. However, for a given comparison and set of genes tested, there appeared to usually only be a substantial drop in the number of genes identified using DESeq1. A description of “Differences between DESeq and edgeR” published in 2013 cited “comparison studies” indicating that there was no single best method [9], and DESeq1 was one of the methods used in our earlier publication [49]. However, this is not a comprehensive review of all possible comparisons with larger sample sizes. Nevertheless, this raises the possibility that some methods may show relative improvement in concordance for differentially expressed gene lists as the sample size increases. In fact, the DESeq2 publication cites an earlier reference that gene filtering can improve statistical power [84], when indicating that DESeq2 omits genes with mean normalized counts below a predefined threshold [14].

For all comparisons, the gene identified to vary at the protein level also varied with |fold-change| > 1.2 and FDR < 0.05 in the RNA-Seq data. A lower fold-change was used with an expectation of higher heterogeneity in patient data, but that might not have been necessary for these comparisons. A plot of the size of the TCGA BRCA differentially expressed gene lists with criteria matching the cell line comparisons (|fold-change| > 1.5 and FDR < 0.05) is provided in **Supplemental Figure S8B**, where more HER2 comparisons identify less than 2000 total differentially expressed genes. If no FPKM filter is applied, then the Wilcoxon test becomes the outlier with a higher gene count (rather than DESeq1 being the outlier with the lower gene count). Additionally, there is an interesting observation that the gene related to the immunohistochemistry strain was high ranked by fold-change (**Figure 4**) as well as some of the p-value calculations (if using limma-trend, ANOVA, or Wilcoxon rank sum).

**Figure 4:**
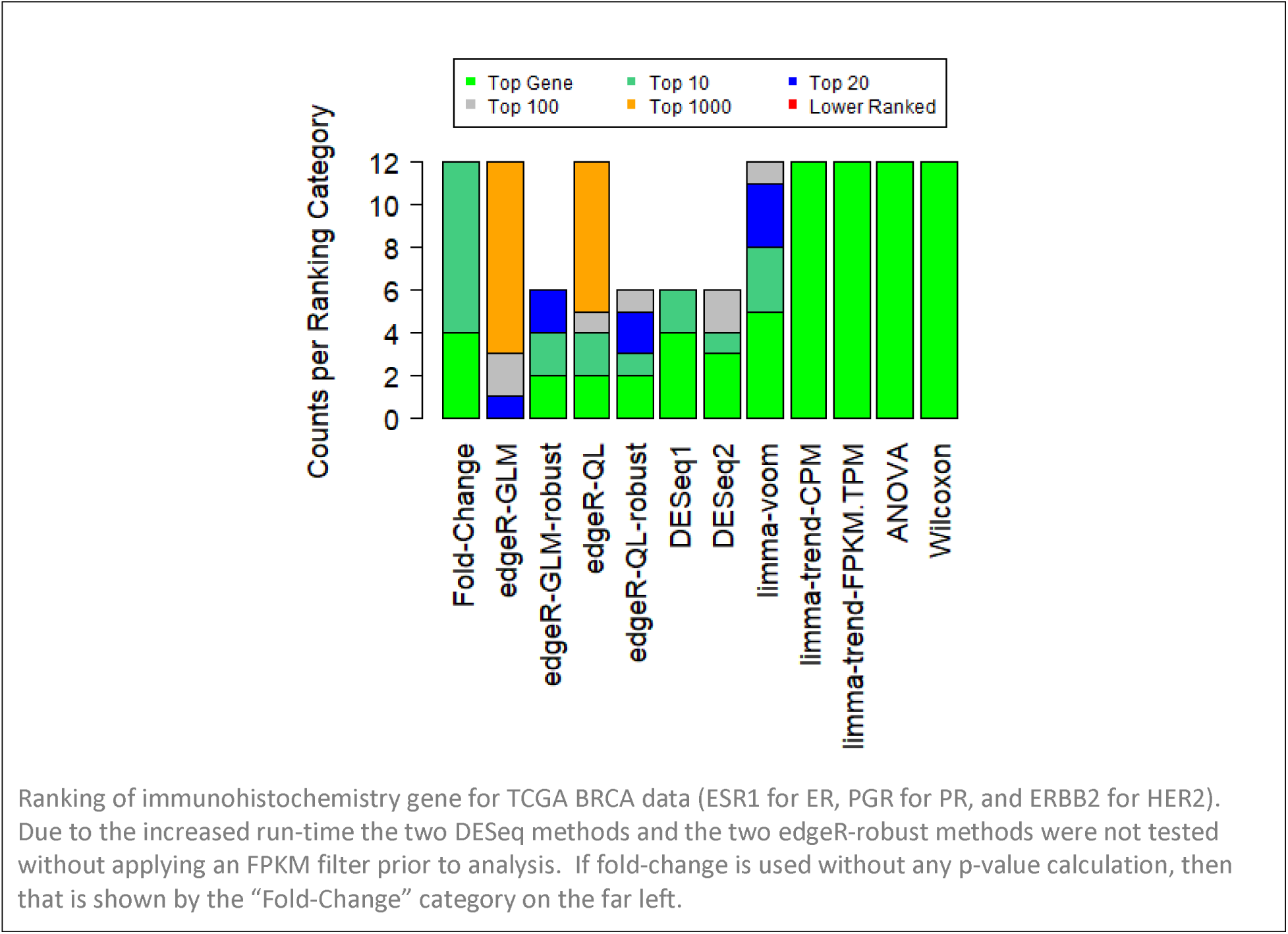
TCGA BRCA RNA-Seq Ranking for Immunohistochemistry Gene

The gene list criteria can be further expanded to only use a threshold of FDR < 0.05, without a threshold (**Supplemental Figure S8C**). This causes some additional divergence, and there is potentially more benefit to applying an FPKM filter prior to differential expression. Additionally, the more conservative nature of the edgeR (GLM) implementation without the robust dispersion estimate can be observed. However, we would typically <u>impose</u> some type of fold-change threshold along with an FDR threshold. For example, the number of genes or transcripts identified can be greater than 30,000. For cell line comparisons, we may try to select 500-1,500 genes (in a given direction of genes): even if a total of 3,000 gene is considered, then a gene list with 10,000 genes is already larger than may be ideal. In (**Supplemental Figure S8A**, focusing on annotations with a larger fraction of coding genes reduces the total count, and this may also reflect more of the genes with annotations for functional enrichment.

At the other end of the ranking categories in **Figure 4**, comparisons where the “robust” dispersion estimate was not used for edgeR also show lower ranking for the immunohistochemistry gene. Thus, the conclusion that the “robust” dispersion estimation can be helpful when running edgeR shows some support in both cell line and patient data.

### Experimental Format for Preprint Content

We encourage **open discussion** through the **Disqus** comment system with this preprint. However, please note that we plan to create a more concise peer-reviewed version. The goal is to reduce the need to have corrections for minor errors. Either way, the plan is to provide full information through the preprint system.

For example, we have created a **<u>Supplemental Results</u>** section. We believe this contains content that may relate to potential reader questions, but this content will likely be removed from a short report or brief communication (such as a potential future submission to a formal peer reviewed journal). The goal is to be more flexible about correcting minor errors, while also being as complete and transparent as possible.

There are currently two main topics contained within the **<u>Supplemental Results</u>**. First, rationale for the study and citation of other previously published studies using templates related to this study are described (**Supplemental Table S3**, [85–92]). Second, a limited number of comparisons on downstream enrichment results is provided (**Supplemental Table S4**, [26, 70, 71, 73]). Content related to the fold-change versus p-value rankings (**Supplemental Table S3-S4**) may also be removed prior to submission for peer review. However, our goal was to maximize content that could be relevant for discussion in the “*v1*” preprint.

## Discussion

For most projects, this manuscript indicates that a differential expression result using only one differential expression method may be acceptable. However, if you assist with analysis of a large enough number of projects, then the introduction of some problem with a lack of flexibility in the analysis strategy may be inevitable. Additionally, we primarily focus on a single step (differential expression of bulk RNA-Seq data). For example, variation in enrichment results can also be studied [93], and we provide some limited results in that respect that might match the expectation that recovery of the causal gene only represents a fraction of the total ways in which the methods could impact the main conclusions of a publication. While all experiences have not currently been published, we also have some experience to indicate a need to critically assess downstream enrichment results [92]. So, if other steps also require critical assessment before publication, then that is also consistent with the conclusion that a strategy that allows variation in methods on a case-by-case basis should help maximize the minimum quality of analysis among all projects supported.

Some differential expression comparisons yielded similar results with the criteria of |fold-change| > 1.5 and FDR < 0.05 with different p-values used to calculate FDR values, but sometimes the number of differentially expressed genes could vary more than 4-fold with the same set of data and a different p-value method used to identify differentially expressed genes. While comparisons with larger sample sizes were limited, the relative performance of methods to identify the immunohistochemistry gene was different than the relative performance of recovery of the causal gene (knock-down or over-expression) in the smaller cell line experiments. For example, the larger TCGA comparisons were more likely to identify each of the three immunohistochemistry genes as exactly the top-ranked gene (if using limma-trend, ANOVA on log2-FPKM values, or a Wilcoxon Rank-Sum test). In most cases, ranking genes by p-value or FDR was comparable to ranking genes by fold-change in the cell line experiments; however, the causal gene was often not precisely the top ranked gene and there was a small fraction of comparisons where the p-value ranking was considerably worse than the fold-change ranking. In the smaller cell line experiments, edgeR-robust or DESeq2 may be a good starting point for 2-group comparisons; however, we believe it is important to have some way to critically assess and revise results in order to try and achieve a minimal level of quality across a large number of projects. Nevertheless, even though all experiences have not been published, the first author has been testing counts of differentially expressed genes defined with edgeR-robust more often for more recent projects (**Supplemental Table S3**).

In theory, the exact reasons why certain methods could perform better under circumstances might be better understood in the future. For example, perhaps some methods perform better with 2-group comparisons versus a multivariate comparison versus one or more continuous variables. Another possibility is that the number of genes affected could affect the performance of different methods. In this study, we provide both single variable and multivariate analysis, but we don’t directly consider a continuous variable (such as time) and we also don’t know the number of affected genes in advance (even though we provide a large range of affected genes among the full set of cell line comparisons. We also expect the number of genes considered can affect the performance of certain sample processing strategies (such as with miRNA-Seq, organisms with smaller genomes / fewer total genes, etc.), but we only re-analyze raw human data in this study. We believe more precise understanding of when to use a given method in advance would be ideal. It is possible that advances in methods may help, but we still strongly believe that the basis for recommendations/suggests must be explainable [91] and such a strategy might be considered “semi-automated” if human critical assessment is needed for final decisions [94]. However, if all possible complications cannot be predicted, then we believe the general idea of expecting rounds of discussion of analysis to be the best strategy in the near future.

We believe the concept of needing methods testing and time for critical assessment for individual projects is not unique to bulk RNA-Seq differential expression [95]. However, we found this to be a study design that was relatively easy to complete with publicly available data. Another example in a different area of genomic research might be that the TCGA GDC workflow applies five methods for somatic variant calls for the same sample ([96], viewed on 6/7/2022). We have also published a study emphasizing the need for raw data and careful critical assessment in an Amplicon-Seq dataset [97]. We do not currently have a formal preprint or peer-reviewed publication, but we expect there is at least similar levels of complications for single cell RNA-Seq data. For example, variation in estimates for the number of cells in a sample can be seen in this GitHub discussion: https://github.com/xnnba1984/Doublet-Detection-Benchmark/issues/4.

In the clinical setting, we understand the benefit of finding solutions that benefit the patient in as few “rounds” as possible as well as avoiding attempts to provide care that unintentionally causes harm. In one sense, we believe it is important to try and help set the right expectations for genomic applications that can be reasonably automated versus those that are more for hypothesis generation. While largely outside the scope of this publication, we might use the term “hypothesis generation” to describe genomic methods that can be helpful but can’t be guaranteed for every individual patient (or project, in a research setting). For example, even without considering the full set of results/goals that could be defined for RNA-Seq data, the threshold for calling an application “hypothesis generation” could arguably apply to the narrow goal of recovering the casual gene with edgeR-robust or DESeq2 in a small size cell line experiment given that the exact thresholds had to be modified from |fold-change| > 1.5 and FDR < 0.05 order to recover the causal gene for all comparisons tested in this study with edgeR-robust (GLM implementation). We also don’t want to underemphasize the effort needed for applications that may be expected to be relatively straightforward over time and between batches. Nevertheless, if limitations are known for research methods, then setting appropriate expectations can be important for the pace of scaling up applications and/or carefully accessing utility for a later clinical use.

Writing and/or modifying code for every project can add human error. If a representative set of results can be reproduced after re-running code to deposit publicly, then we hope that can help with the debugging process. We also hope that reviewers ask questions about method selection and troubleshooting during the peer review process, both in terms of literature familiarity as well as first-hand experience with the data specifically presented in the publication. We believe that use of an analysis “template” helps reduce human error, but it is certainly not 100% effective by itself. We acknowledge that those strategies may not provide the absolute best possible solution, and we certainly encourage further discussion on this topic.

Nevertheless, we believe that the concept of providing multiple rounds of analysis and discussion for “initial” results that may require additional modification at later dates in a research setting is important for project planning for bulk RNA-Seq data, and we have provided a number of results to support that conclusion. We also hope that this publication helps with setting the right expectations for new collaborations in providing shared analysis support.

Again, we encourage **open discussion** through the **Disqus** comment system with this preprint.

## Supporting information

Supplemental Results

## Acknowledgements

There are a very large number of labs and individuals that should be acknowledged for the RNA-Seq gene expression template used for core support. When they provided agreement for a public acknowledgement, they are listed within the README.md file: https://github.com/cwarden45/RNAseq_templates/blob/master/TopHat_Workflow/README.md. More recently, we would also like to thank Jose Enrique Montero Casimiro for valuable input for result summaries. There have also been newer projects supported with code derived from those templates, and those labs are acknowledged on the Zenodo page (https://zenodo.org/records/3378055).

Additionally, we would like to thank Ching Ouyang for a journal discussion related to edgeR implementations, which influenced the experimental design for part of this study (as well as presenting the reference for the non-parametric Wilcoxon test for RNA-Seq data). In the interests of open and transparent research with communication of gradually increasing formality, the ideas for this manuscript have also been presented in the following blog post: https://cdwscience.blogspot.com/2019/11/requiring-at-least-some-methods-testing.html

## Funding

The Integrative Genomics Core is supported by the National Cancer Institute Comprehensive Cancer Center grant (P30CA033572). The content is solely the responsibility of the authors and does not necessarily represent the official views of the National Institutes of Health.

## Author Contributions

C.D.W performed the analysis, designed the study, and wrote the paper. X.W. provided supervision and wrote the paper.

## Conflict of Interest

The authors declare no conflict of interest.

## Notes

https://zenodo.org/records/3378055

https://sourceforge.net/projects/rnaseq-deg-methodlimit/

### Competing Interest Statement

The authors have declared no competing interest.

