## Supplemental Results for "Critical Differential Expression Assessment for Individual Bulk RNA-Seq Projects"

***Additional Evidence for Value in Some Methods Testing for Every Project***

The original basis for putting together systematic analysis of public data was the overall experiences when using a template for RNA-Seq analysis to support various labs, and the public data in this study could arguably be considered validation data for an observation across individual lab’s projects. All analysis provided to labs does not have an associated publication, and the methods testing is usually not directly described in the associated paper. Nevertheless, the observation that different methods are used in different papers (Dalasanur Nagaprashantha et al. 2018; Petrossian et al. 2018; Oh et al. 2018; Kanaya et al. 2019; Ubina et al. 2019; Merz et al. 2022; Su et al. 2019; Guo et al. 2023) can be reported as indirect evidence for the conclusion from this publication (**Supplemental Table S3**). One complication is that the template itself evolved over time, largely due to feedback from the collaborating labs. Additionally, tor the “initial” results, there was also some subjective assessment about what the best differential expression method might be to try first (based upon factors like the number of genes identified, the symmetry of up-regulated versus down-regulated genes, and visual inspection of heatmaps created using an independently calculated gene expression value for a candidate method). Thus, all factors considered are not described in this paper, and quantifying the contribution of all factors with the studies that provide appropriate experimental control conditions may sometimes be difficult. Nevertheless, providing some methods testing for every project is not just a theoretical recommendation, but it is something that has been put into practice. DESeq2, edgeR, and limma-voom were used as the main result for at least one publication, and you can occasionally see multiple methods were used within the same study for different comparisons (Kanaya et al. 2019; Merz et al. 2022).

***Limited Assessment of Downstream Enrichment for Selected Datasets***

Limitations in gene signatures for selected transcriptional regulators were indirectly described a previous study showing that it is sometimes helpful or necessary to define and apply custom gene signatures (Warden et al. 2013), but those did not include any of the causal genes in this study. So, we can test a modified version of the previously published analysis for selected comparisons where similar public gene sets exist for the causal knock-out or over-expression genes (for DESeq2 and edgeR-robust-GLM results). As described in the Methods, GSEA and BD-Func analysis was performed in multiple ways for each gene set, and Enrichr was also tested for over-representation of gene sets for causal genes in this study.

For BMI1, the E-MTAB-4237 gene lists and/or normalized expression could be analyzed with GSEA, Enrichr, and BD-Func for c6 MSigDB signatures (and GSEA and BD-Func were used for c2 MSigDB signatures, **Supplemental Table S4**). The c6 gene sets (BMI1_DN.V1_UP, BMI1_DN.V1_DN, BMI1_DN_MEL18_DN.V1_UP, BMI1_DN_MEL18_DN.V1_DN) showed a strong trend for 2 or 3 methods, but in the opposite direction as would be expected. As described in the methods, re-analysis of the original microarray dataset (GSE7578 (Wiederschain et al. 2007)) for the c6 signatures yields similar results. While the rankings were not especially high, there were results with an FDR < 0.25 (and some with FDR < 0.05) based upon the c2 signatures (DOUGLAS_BMI1_TARGETS_UP and DOUGLAS_BMI1_TARGETS_DN).

For HIF1A, the publication associated with E-MTAB-1994 (Choudhry et al. 2014) references using the same protocol as the publication for the MSigDB signatures (Elvidge et al. 2006). There are also similar experimental conditions (such as 21% oxygen for normoxia and 1% oxygen for hypoxia in MCF7 cells), as well as having an overlapping author. At the same time, the technology used for measuring gene expression is different between the publications (a microarray signature is being applied to RNA-Seq data). So, there may be some amount of over-fitting and/or more similar results than a more independent experiment. However, arguably, this might mean that these signatures serve an even stronger positive control. If considered in that way, it might be encouraging that ELVIDGE_HIF1A_TARGETS_UP was ranked first when using GSEA with pre-ranked scores based upon signed log-transformed DESeq2 or edgeR-robust-GLM p-values (**Supplemental Table S4**). Those same scores also yielded highly significant BD-Func results, although the ranking was somewhat lower.

For GATA6, the ZHANG_GATA6_TARGETS_UP and ZHANG_GATA6_TARGETS_DN signatures did not show good performance on the E-MTAB-6756 data (**Supplemental Table S4**).

We consider estimations of frequency for difficulties in downstream analysis to be outside of the scope of this study. These limited examples illustrate complications that can occur with downstream analysis. Taken together with previous results for gene signatures for different causal genes (Warden et al. 2013), this provides some corroboration to the expectation that recovery of the causal gene in differentially expressed gene lists only comprises a fraction of the total ways in which hypotheses generated by “initial” RNA-Seq analysis throughout the entire workflow need to be carefully evaluated.

Dalasanur Nagaprashantha, Lokesh, Jyotsana Singhal, Hongzhi Li, Charles Warden, Xueli Liu, David Horne, Sanjay Awasthi, Ravi Salgia, and Sharad S. Singhal. 2018. '2&#x0027;-Hydroxyflavanone effectively targets RLIP76-mediated drug transport and regulates critical signaling networks in breast cancer', *Oncotarget*, 9.

Elvidge, Gareth P., Louisa Glenny, Rebecca J. Appelhoff, Peter J. Ratcliffe, Jiannis Ragoussis, and Jonathan M. Gleadle. 2006. 'Concordant Regulation of Gene Expression by Hypoxia and 2-Oxoglutarate-dependent Dioxygenase Inhibition: THE ROLE OF HIF-1α, HIF-2α, AND OTHER PATHWAYS*', *Journal of Biological Chemistry*, 281: 15215-26.

Guo, Linlin, Atish Mohanty, Sharad Singhal, Saumya Srivastava, Arin Nam, Charles Warden, Sravani Ramisetty, Yate-Ching Yuan, Hyejin Cho, Xiwei Wu, Aimin Li, Manik Vohra, Srinivas Vinod Saladi, Deric Wheeler, Leonidas Arvanitis, Erminia Massarelli, Prakash Kulkarni, Yiming Zeng, and Ravi Salgia. 2023. 'Targeting ITGB4/SOX2-driven lung cancer stem cells using proteasome inhibitors', *iScience*, 26: 107302.

Kanaya, Noriko, Lauren Bernal, Gregory Chang, Takuro Yamamoto, Duc Nguyen, Yuan-Zhong Wang, June-Soo Park, Charles Warden, Jinhui Wang, Xiwei Wu, Timothy Synold, Michele Rakoff, Susan L Neuhausen, and Shiuan Chen. 2019. 'Molecular Mechanisms of Polybrominated Diphenyl Ethers (BDE-47, BDE-100, and BDE-153) in Human Breast Cancer Cells and Patient-Derived Xenografts', *Toxicological Sciences*, 169: 380-98.

Merz, Karla E., Ragadeepthi Tunduguru, Miwon Ahn, Vishal A. Salunkhe, Rajakrishnan Veluthakal, Jinhee Hwang, Supriyo Bhattacharya, Erika M. McCown, Pablo A. Garcia, Chunxue Zhou, Eunjin Oh, Stephanie M. Yoder, Jeffrey S. Elmendorf, and Debbie C. Thurmond. 2022. 'Changes in Skeletal Muscle PAK1 Levels Regulate Tissue Crosstalk to Impact Whole Body Glucose Homeostasis', *Frontiers in Endocrinology*, 13.

Oh, Eunjin, Miwon Ahn, Solomon Afelik, Thomas C. Becker, Bart O. Roep, and Debbie C. Thurmond. 2018. 'Syntaxin 4 Expression in Pancreatic β-Cells Promotes Islet Function and Protects Functional β-Cell Mass', *Diabetes*, 67: 2626-39.

Petrossian, Karineh, Noriko Kanaya, Chiao Lo, Pei-Yin Hsu, Duc Nguyen, Lixin Yang, Lu Yang, Charles Warden, Xiwei Wu, Raju Pillai, Lauren Bernal, Chiun-Sheng Huang, Laura Kruper, Yuan Yuan, George Somlo, Joanne Mortimer, and Shiuan Chen. 2018. 'ERα-mediated cell cycle progression is an important requisite for CDK4/6 inhibitor response in HR+ breast cancer', *Oncotarget*, 9.

Su, Yapeng, Xiang Lu, Guideng Li, Chunmei Liu, Yan Kong, Jihoon W. Lee, Rachel Ng, Stephanie Wong, Lidia Robert, Charles Warden, Victoria Liu, Jie Chen, Zhuo Wang, Yezi Yang, Hanjun Cheng, Alphonsus H. C. Ng, Guangrong Qin, Songming Peng, Min Xue, Dazy Johnson, Yu Xu, Jinhui Wang, Xiwei Wu, Ilya Shmulevich, Qihui Shi, Raphael Levine, Antoni Ribas, David Baltimore, Jun Guo, James R. Heath, and Wei Wei. 2019. 'Kinetic Inference Resolves Epigenetic Mechanism of Drug Resistance in Melanoma', *bioRxiv*: 724740.

Ubina, Teresa, Martha Magallanes, Saumya Srivastava, Charles D. Warden, Jiing-Kuan Yee, and Paul M. Salvaterra. 2019. 'A Human Embryonic Stem Cell Model of Aβ-Dependent Chronic Progressive Neurodegeneration', *Frontiers in Neuroscience*, 13.

Warden, Charles D., Noriko Kanaya, Shiuan Chen, and Yate-Ching Yuan. 2013. 'BD-Func: a streamlined algorithm for predicting activation and inhibition of pathways', *PeerJ*, 1: e159.

Wiederschain, Dmitri, Lin Chen, Brett Johnson, Kimberly Bettano, Dowdy Jackson, John Taraszka, Y. Karen Wang, Michael D. Jones, Michael Morrissey, James Deeds, Rebecca Mosher, Paul Fordjour, Christoph Lengauer, and John D. Benson. 2007. 'Contribution of Polycomb Homologues Bmi-1 and Mel-18 to Medulloblastoma Pathogenesis', *Molecular and Cellular Biology*, 27: 4968-79.
