## Supplementary figures and images for "Critical Differential Expression Assessment for Individual Bulk RNA-Seq Projects"

### Supplemental Figure S1

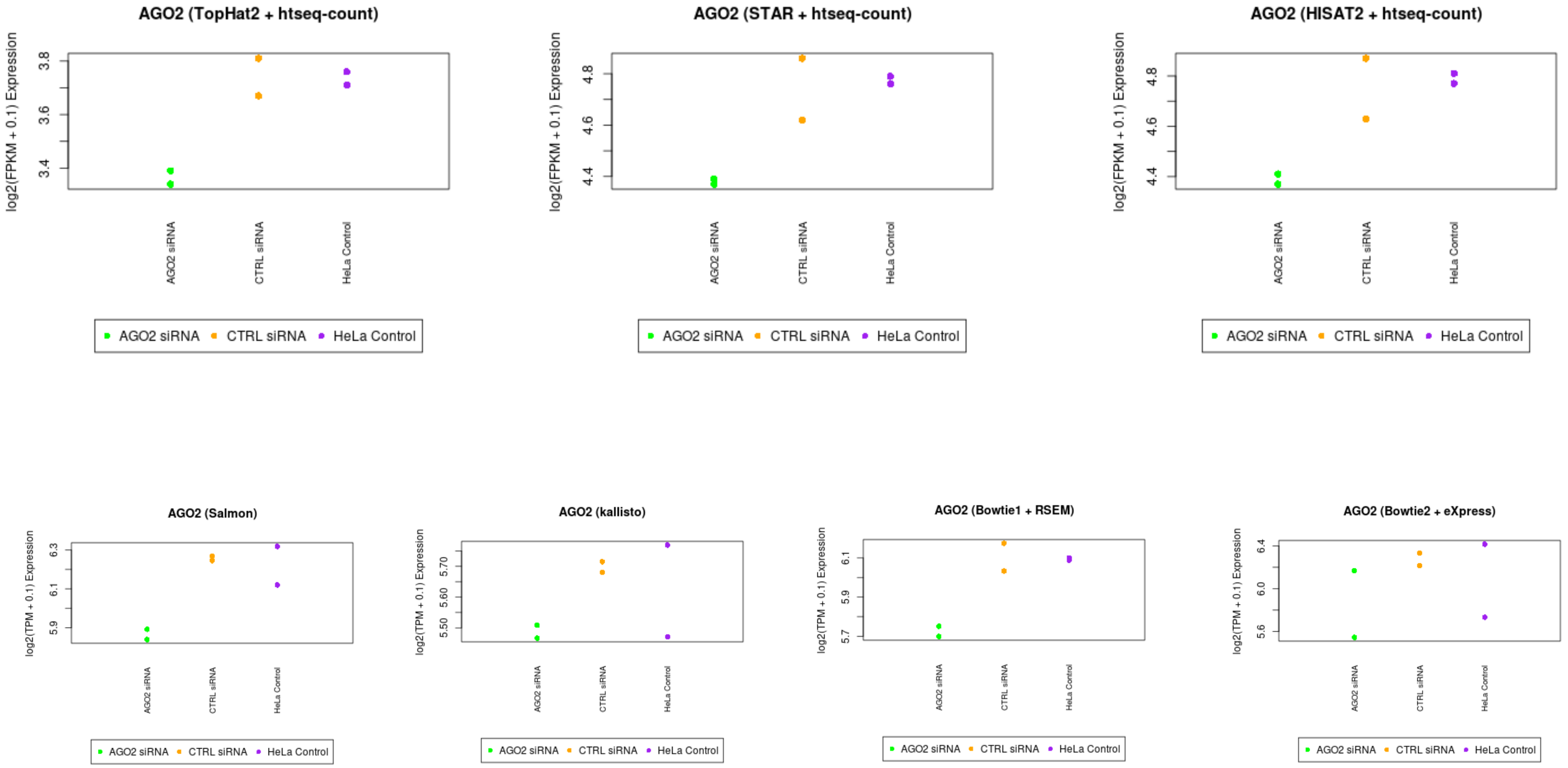

### Supplemental Figure S2

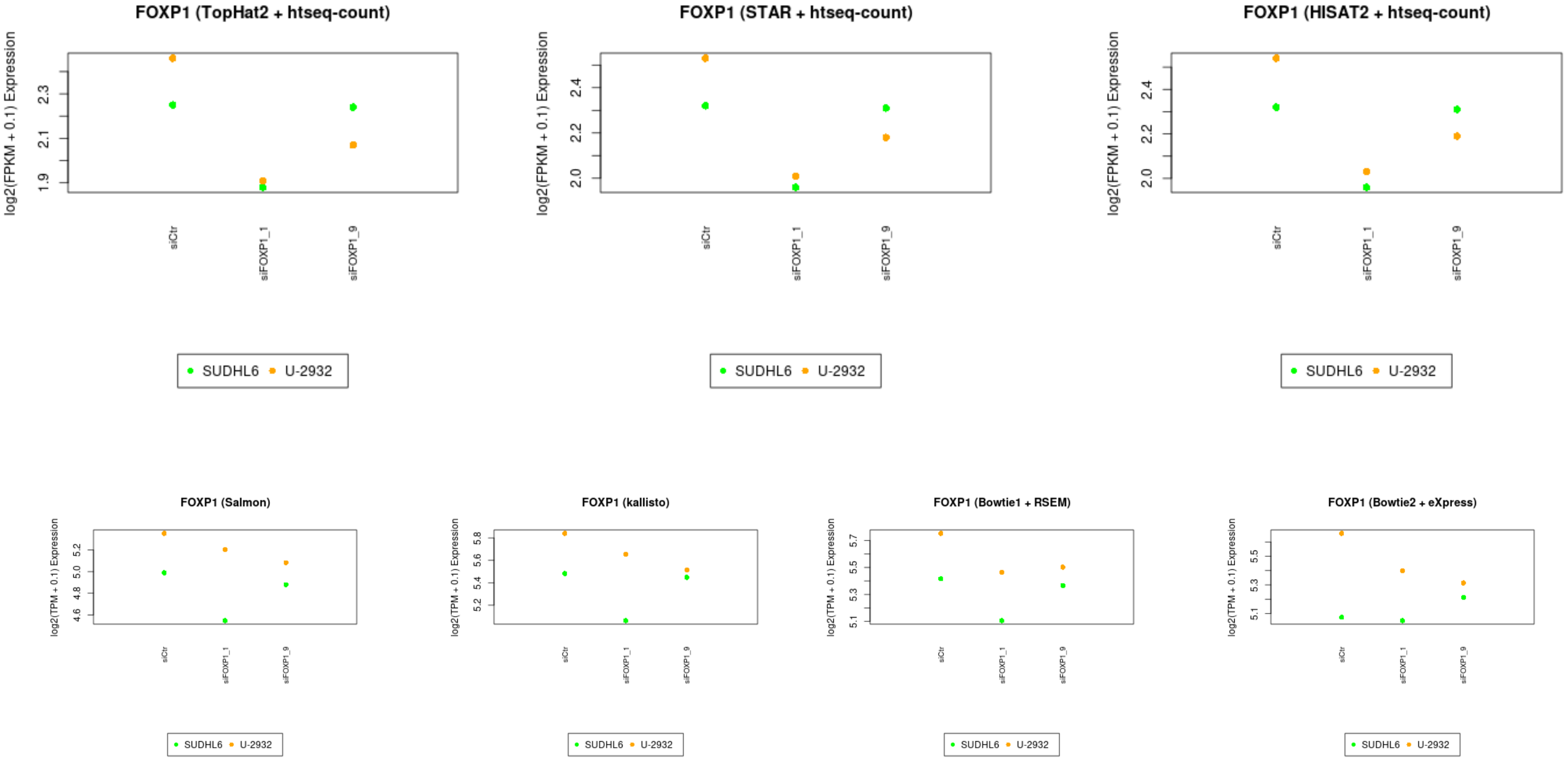

### Supplemental Figure S3

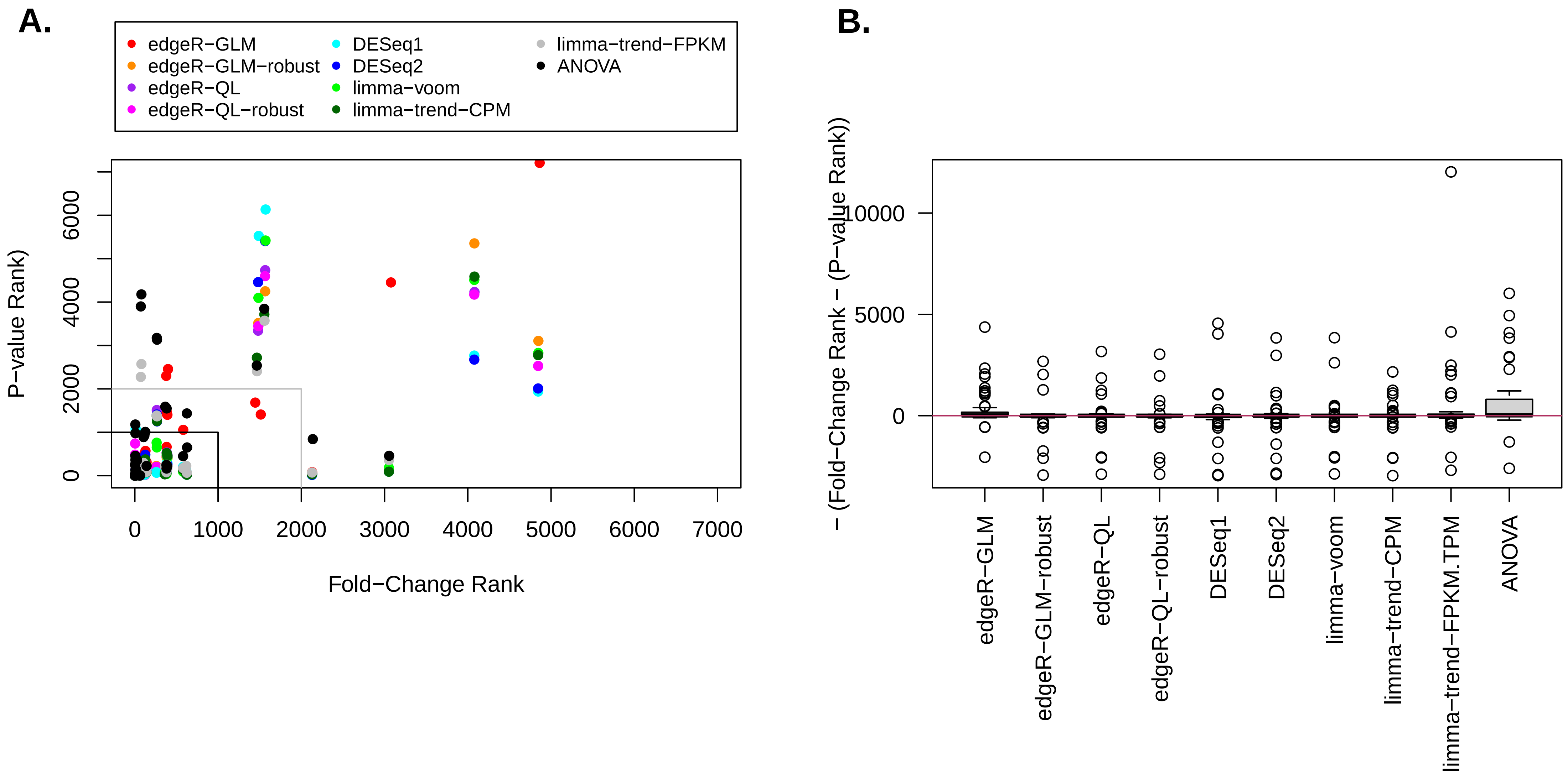

### Supplemental Figure S4

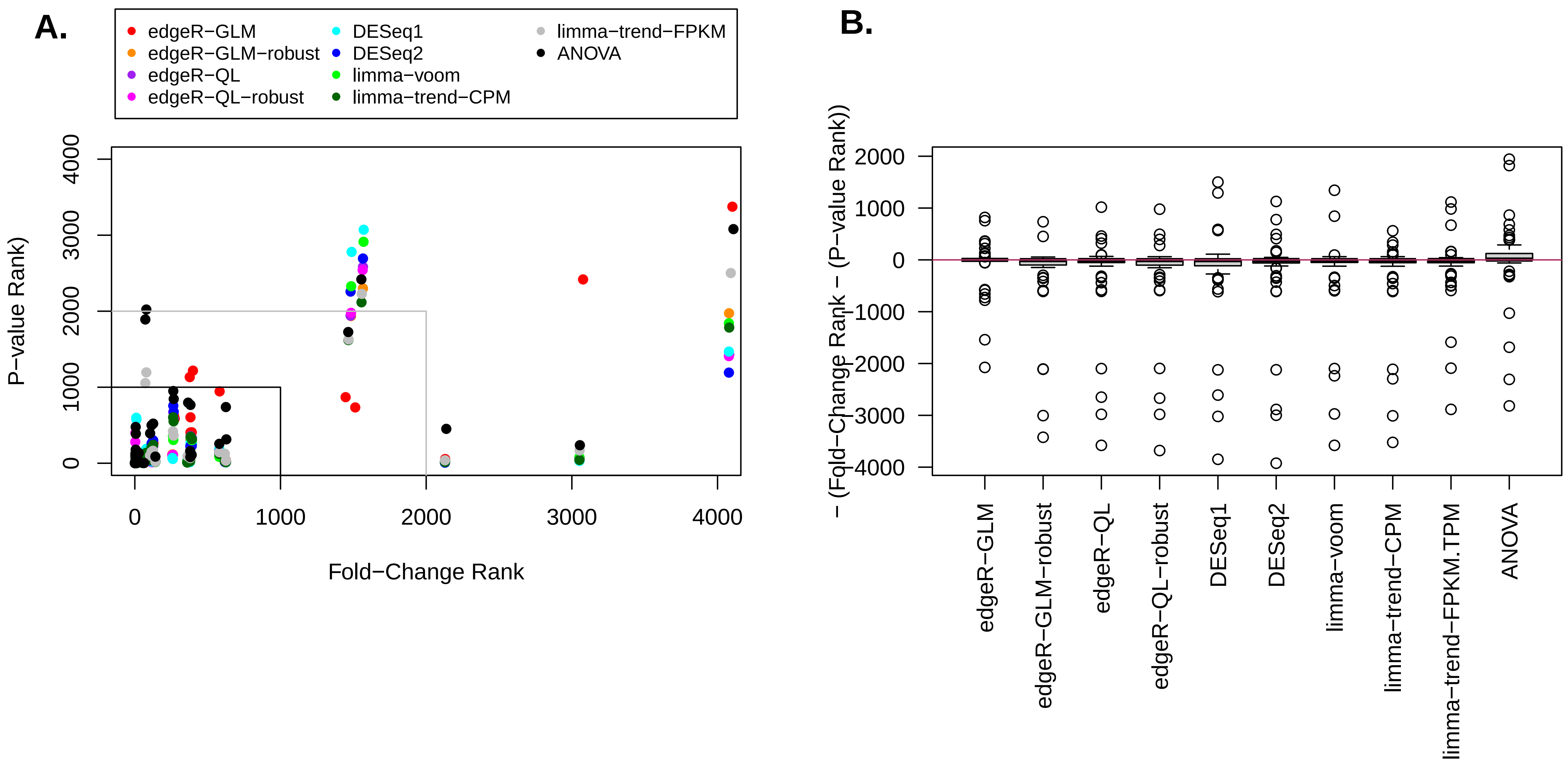

### Supplemental Table S5

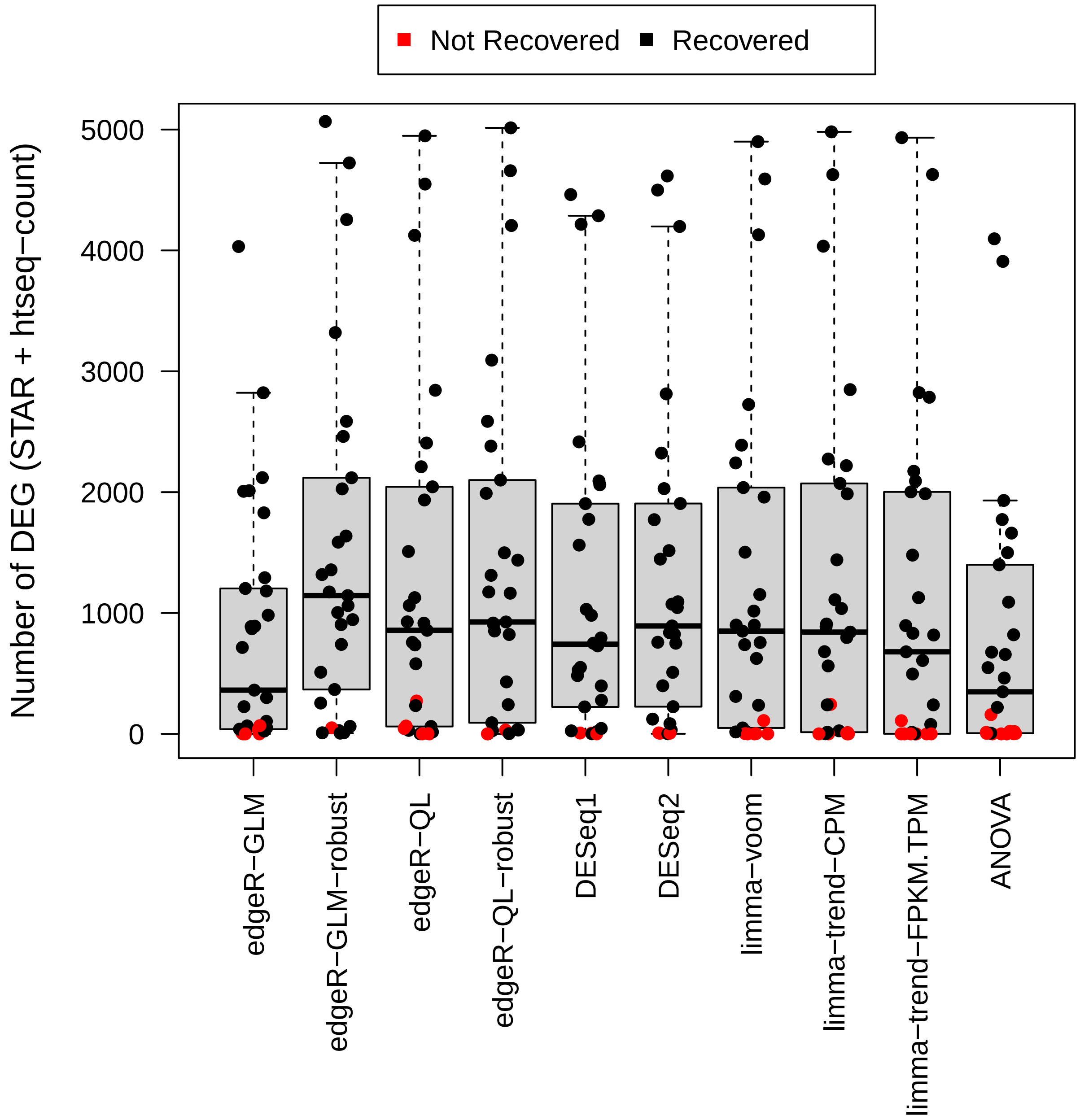

### Supplemental Table S6

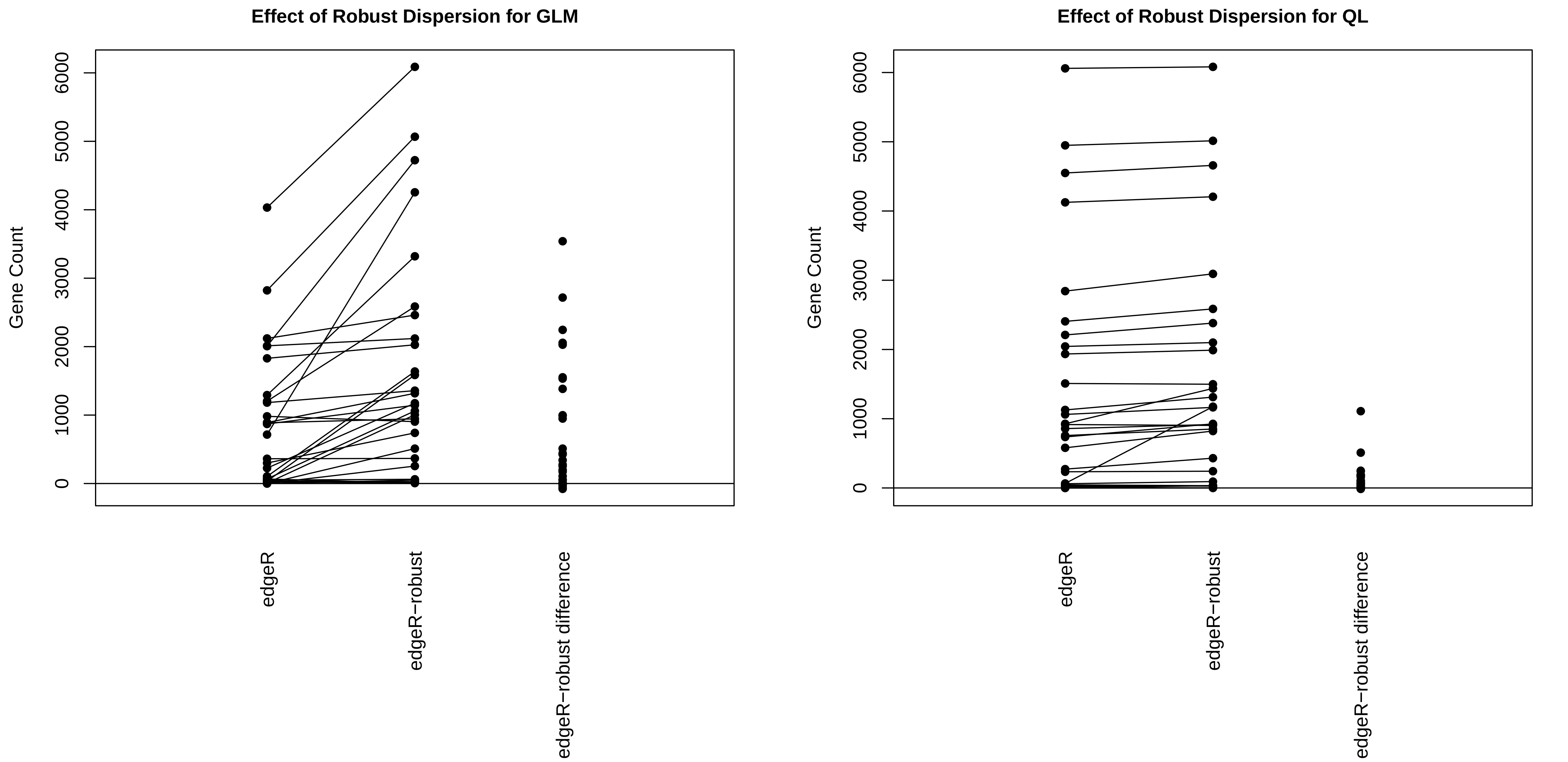

### Supplemental Table S7

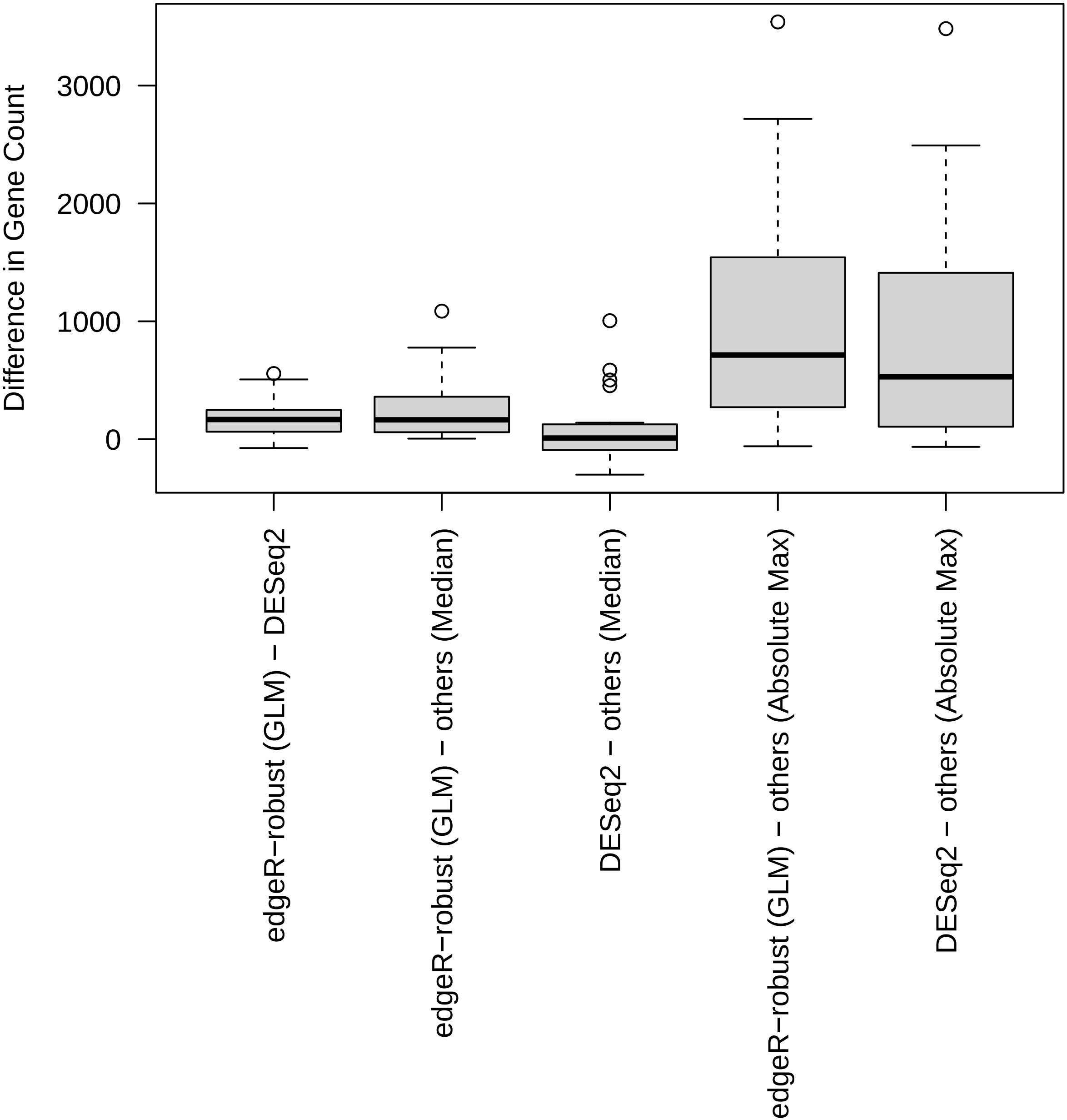

### Supplemental Table S8

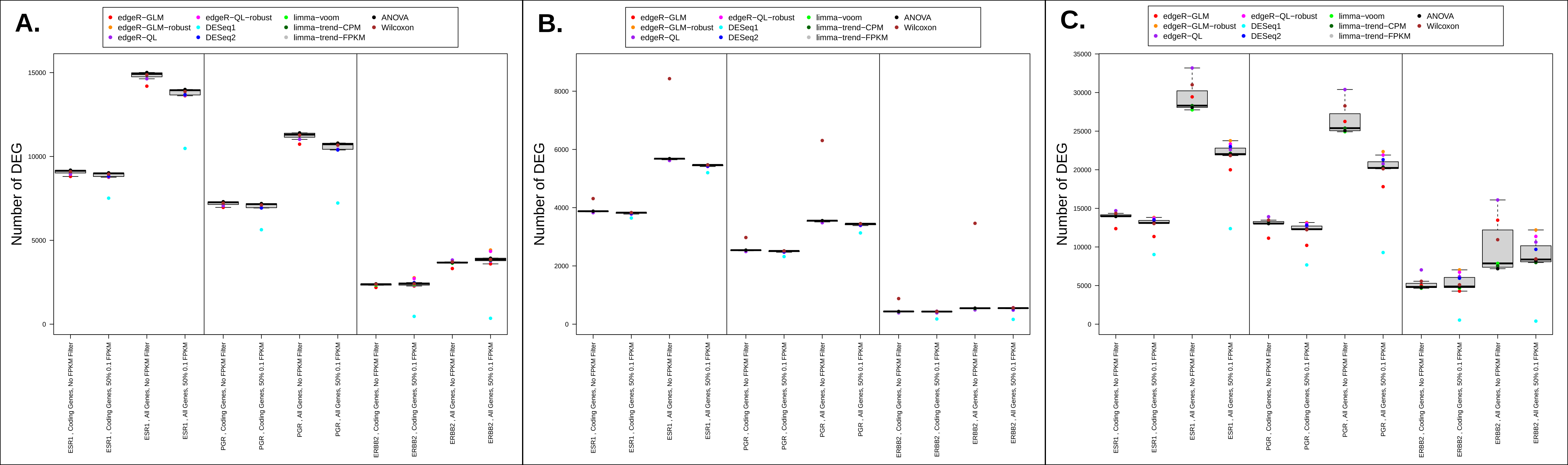
